# Striatal activity during contextual word learning is influenced by children’s reading ability

**DOI:** 10.64898/2026.07.06.736136

**Authors:** Nilgoun Bahar, Desislava Arabadzhiyska, Hannah Jones, Sonia Singh, Miykael Davis, Jessie Ricketts, Pablo Ripollés, Saloni Krishnan

## Abstract

Contextual word learning is a fundamental mechanism for vocabulary acquisition during childhood. In adults, successful inference of word meaning from context is intrinsically rewarding, and is associated with greater enjoyment and greater activity in reward-related brain regions. Whether similar reward mechanisms support word learning in children, and whether they differ as a function of ability, remains unknown. We used functional magnetic resonance imaging (fMRI) to examine neural responses during contextual word learning in 25 children aged 11-13 years with typical reading skills and in 20 age-matched children with dyslexia. Neurotypical readers showed enhanced activation in core reward-processing regions, including the ventral striatum, when successfully learning the meanings of novel words. In contrast, children with dyslexia did not exhibit comparable reward-related responses despite performing the same task. Crucially, this group difference was specific to word learning, as no significant group differences were observed in ventral striatal responses during a non-linguistic monetary reward task. In addition, to confirm the behavioural relevance of these neural findings, we examined an age-matched, independent sample of children. We found that stronger reading skills were associated with greater enjoyment during successful word learning. Together, these results suggest that interactions between reward and language systems during contextual word learning is influenced by reading proficiency. Reduced intrinsic reward responses to successful language learning may contribute to differences in reading development and have implications for the design of more engaging and effective reading interventions for struggling readers.

**Significance statement:** We investigated how school-age children’s brains learn new words from context, highlighting the important role of reward systems in this process. We show that ventral striatum, a core region associated with reward processing, is activated when children learn the meaning of new words. This suggests learning the meaning of a new word can be its own reward, driving further learning and engagement. Yet, children with dyslexia showed reduced brain activity in the ventral striatum during language learning, even when they successfully learned words. These findings underscore the importance of understanding the mechanisms through which learning becomes rewarding, with implications for designing more engaging and effective reading interventions

## 1. Introduction

Learning can be its own reward. Acquiring new information engages the brain’s dopaminergic reward circuitry, including the ventral striatum and ventral tegmental area (VTA), and this activation promotes long-term potentiation and memory consolidation (Bromberg-Martin et al., 2010; Lisman & Grace, 2005; Wittmann et al., 2016). Learning the meaning of a new word similarly recruits core reward-processing circuits; in adults, successful acquisition of novel word meanings from context is associated with activation of the ventral striatum and VTA (Ripollés et al., 2014, 2016) and pharmacological manipulation of dopamine provides causal evidence for the involvement of this system in word learning (Ripollés et al., 2018). These findings together suggest that dopaminergic reward systems contribute to successful word learning and fuel memory for words. An implicit assumption of existing studies is that these intrinsic reward responses during learning are experienced uniformly across development and across individuals. In this study, we test this assumption, assessing whether the recruitment of reward systems during word learning is similar in children with varying levels of reading proficiency.

The experience of intrinsic reward in word learning is typically investigated using contextual word learning, in which readers infer word meanings from the available context. This naturalistic process is a major source of vocabulary growth in both children and adults (Nagy et al., 1985; Nation, 2017; Swanborn & De Glopper, 1999). By approximately eight years of age, children can acquire new word meanings from written context without explicit instruction (Kuhn & Stahl, 1998; Nagy et al., 1985). In adults, successful contextual word learning is rated as enjoyable even in the absence of external feedback or incentives and after controlling for factors such as novelty and effort (Angwin et al., 2019; Ripollés et al., 2014; Zaka et al., 2026). Similarly, children and adolescents aged 10-18 years report enjoyment following successful inference of word meanings from context (Bains et al., 2024). These findings suggest that intrinsic reward may support vocabulary acquisition across development. However, whether children recruit the same neural reward circuitry as adults during word learning, and whether this engagement is modulated by reading ability, remain unknown.

Reward responsivity varies as a function of experience and environmental context. For example, children from lower socioeconomic backgrounds and those who have experienced adverse childhood events exhibit reduced reward responsivity (Decker et al., 2024; Lloyd et al., 2022). In the domain of reading, ability influences motivation: children with dyslexia read less frequently than their neurotypical peers (Siegel, 2006; Stanovich, 1986) and often report reduced reading motivation (Jones et al., 2025; Van Bergen et al., 2018). Such differences in reading experience may influence how reward systems are engaged during learning. During contextual word learning, learners must internally monitor their learning success, evaluating whether they have accurately inferred a word’s meaning without explicit feedback (Caras et al., 2022; Ripollés et al., 2016). Stronger readers with more reading experience may have greater insight into their learning accuracy and performance and therefore may be more likely to experience intrinsic reward. Whether children with dyslexia experience comparable reward-related neural responses during word learning remains unknown.

Lastly, understanding individual differences in the experience of reward is important because dopaminergic signalling enhances not only memory formation but also motivation to pursue future rewards (Bromberg-Martin et al., 2010; Gruber & Ranganath, 2019). Successful learning may therefore foster a positive feedback loop that increases engagement with future learning opportunities. In the present study, we adapted a previously validated contextual word learning paradigm (Ripollés et al., 2014) for use with 53 children aged 11-13 years, approximately half of whom had a diagnosis of dyslexia, while acquiring fMRI data. Based on prior behavioural findings (Bains et al., 2024) and neural evidence in adults (Ripollés et al., 2014, 2016, 2018), we hypothesised that successful word learning would elicit activation in reward-related neural circuitry in children and that the magnitude of this response would vary as a function of children’s reading ability.

## 2. Materials and Methods

We report data from two studies, a new fMRI study (section 2.1) and a secondary analysis of an existing behavioural study (section 2.2).

### 2.1 fMRI study of contextual word learning

#### 2.1.1 Ethics

Ethical approval for this study was received from Royal Holloway, University of London’s ethical review committee (Project Code: 1849). Prior to participation, we obtained written informed consent from parents/guardians and written assent from children.

#### 2.1.2 Participants and study criteria

We recruited 53 children aged 11-13 years from local schools, social media advertisements, and organisations supporting individuals with reading difficulties (e.g., the British Dyslexia Association). Twenty-eight children had typical language and reading development (typically developing [TD] group; *M* age = 12.5 years, *SD* = 0.58; 15 males), and 25 had a formal diagnosis of dyslexia or were receiving support for reading difficulties (RD group; *M* age = 12.4 years, *SD* = 0.83; 13 males).

Inclusion criteria were: (1) English as the primary language spoken from early childhood in the United Kingdom and (2) a nonverbal reasoning standard score above 70 on the Matrix Reasoning subtest of the Wechsler Intelligence Scale for Children–Fourth Edition (WISC-IV; Wechsler, 2004). Exclusion criteria included: (1) a diagnosis of attention-deficit/hyperactivity disorder, autism spectrum disorder, or another known neurodevelopmental or genetic condition (e.g., Down syndrome); (2) a history of neurological symptoms such as seizures; (3) a score >7 on the Hyperactivity subscale of the Strengths and Difficulties Questionnaire (SDQ; Goodman, 1997); (4) a score >15 on the Social Communication Questionnaire (SCQ; Rutter et al., 2003); or (5) any contraindication to MRI.

Data from one participant were excluded because the scan was not completed. Data from an additional seven participants were excluded because of excessive head motion during MRI acquisition. The final sample comprised 25 TD children and 20 children in the RD group. Table 1 in the next section shows the demographic details and neuropsychological results from the final sample (*N* = 45). The two groups were closely matched on neuropsychological scores, including nonverbal reasoning, and differed primarily in their decoding abilities.

**Table 1.** Demographic and neuropsychological results.

|  | **TD**<br>**(*n* = 25)** | **RD**<br>**(*n* = 20)** | **p-value** | **Group**<br>**differences** |
| --- | --- | --- | --- | --- |
| **Demographics** |  |  |  |  |
| Mean age in years | 12.45 (0.6) | 12.45 (0.9) | 0.982 | --- |
| Age range | 11;0 – 13;10 | 11;6 – 13;9 | --- | --- |
| Sex (Male:Female) | 14:11 | 11:9 | 1.000 | --- |
| **Neuropsychological test battery** |  |  |  |  |
| WISC-IV |  |  |  |  |
| Matrix Reasoning <sup>1</sup> | 9.8 (3.1) | 9.6 (3.6) | 0.776 | --- |
| ROWPVT-4 | 116.4 (8.6) | 112.8 (22.4) | 0.509 | --- |
| CELF-V |  |  |  |  |
| Recalling Sentences <sup>1</sup> | 10.9 (2.6) | 8.9 (3.6) | 0.045 | RD < TD |
| TOWRE-2 |  |  |  |  |
| Sight Word Efficiency | 94.6 (11.7) | 81.7 (17.5) | **0.007** | RD < TD |
| Phonemic Decoding Efficiency | 101.1 (12.6) | 80.1 (14.4) | **<0.001** | RD < TD |
| YARC |  |  |  |  |
| Reading Rate | 109.8 (11.4) | 95.3 (19.9) | **0.007** | RD < TD |
| Reading Comprehension | 111 (11.2) | 106.8 (13.2) | 0.265 | RD < TD |
| **Questionnaires** |  |  |  |  |
| MRP <sup>2</sup> | 21.3 (3.7) | 21.1 (3.8) | 0.873 | --- |
| AMI-CV <sup>3</sup> | 20 (6.3) | 20.4 (7.1) | 0.859 | --- |
| **Word learning paradigm** |  |  |  |  |
| In-scan Word Recognition (%) <sup>4</sup> | 68.6 (13.1) | 52.2 (18.3) | **0.001** | RD < TD |
| Post-scan Word Recognition (%)<sup>4</sup> | 69.2 (14.6) | 51.8 (13.7) | **<0.001** | RD < TD |

*Note.* Values are reported as mean (*SD*) unless otherwise indicated. The final two columns report statistical comparisons between groups and associated p-values. Significant group differences after Bonferroni correction ( $\alpha = 0.008$ ) are shown in bold. Tests without a superscript denote standard scores ( $M = 100$ , $SD = 15$ ). <sup>1</sup> Scaled scores ( $M = 10$ , $SD = 3$ ). <sup>2</sup> Raw score (maximum = 32). <sup>3</sup> Raw score (maximum = 48). <sup>4</sup> Percent correct (chance performance = 33%). TD = typically developing, RD = dyslexia.

#### 2.1.3 Neuropsychological assessment

We characterised children’s reading, language, cognitive ability, and motivation to read using a battery of standardised tasks and questionnaires, which took approximately 1½ hours. Reading fluency was tested using the Phonological Decoding and Sight Word Reading Efficiency subtests from Test of Word Reading Efficiency-Second Edition (TOWRE-2; Torgesen et al., 2012). The Passage Reading subtest from the York Assessment of Reading Comprehension (YARC; Snowling et al., 2009) was administered to assess reading comprehension. Receptive vocabulary was measured using the Receptive One-Word Picture Vocabulary-4th Edition (ROWPVT-4; Martin & Brownell, 2011b). Expressive grammar was assessed using the Recalling Sentences subtest of the Clinical Evaluation of Language Fundamentals-5th Edition (CELF-V; Wiig et al., 2013). The Matrix Reasoning subtest from the Weschler Intelligence Scale for Children-4^th^ edition (WISC-IV; Wechsler, 2004) was administered as a measure of nonverbal reasoning skills.

Children also completed two questionnaires to assess motivation: the Apathy Motivation Index-Child Version (AMI-CV; Hewitt et al., 2023) and an adapted version of the Motivation to Read Profile (MRP; Gambrell et al., 1996) with eight items indexing self-concept as a reader and value placed on reading. Higher AMI-CV scores indicate greater apathy (i.e., lower motivation), whereas higher scores on our modified MRP questionnaire denote a more positive self-concept as a reader and higher value placed on reading.

#### 2.1.4 Experimental design

##### 2.1.4.1 Contextual word learning task design

Participants completed a child-friendly adaptation of the contextual word learning paradigm developed by Ripollés et al. (2014). They were instructed to learn the meanings of 36 novel “words” (i.e., pseudowords) embedded within pairs of sentences. Stimuli comprised 36 sentence pairs (5-7 words per sentence) ending with a six-letter pseudoword that replaced a noun. All pseudowords conformed to English phonotactic constraints.

Sentence pairs were constructed with an increasing degree of contextual constraint. The first sentence provided limited semantic information (mean cloze probability = 18.84%, *SD* = 16.98%; e.g., “Few countries are now ruled by a *traper*”), whereas the second sentence provided much greater contextual constraint (mean cloze probability = 74.59%, *SD* = 14.08%; e.g., “In the palace lives the king and *traper*”). Participants were expected to infer the meaning of each pseudoword from the combined context of the two sentences (e.g., *traper* = *queen*). These stimuli were previously validated in a developmental sample of English-speaking children and adolescents (Bains et al., 2024; see Zaka et al., 2026 for information on cloze probability determination).

To control for visual processing demands, participants also viewed non-readable control stimuli created by transposing letter strings to produce undecipherable sentences matched in length to the meaningful sentences (Fig. 1A, bottom row). Prior to scanning, participants completed a brief practice session to familiarise themselves with the task.

**Figure 1.**
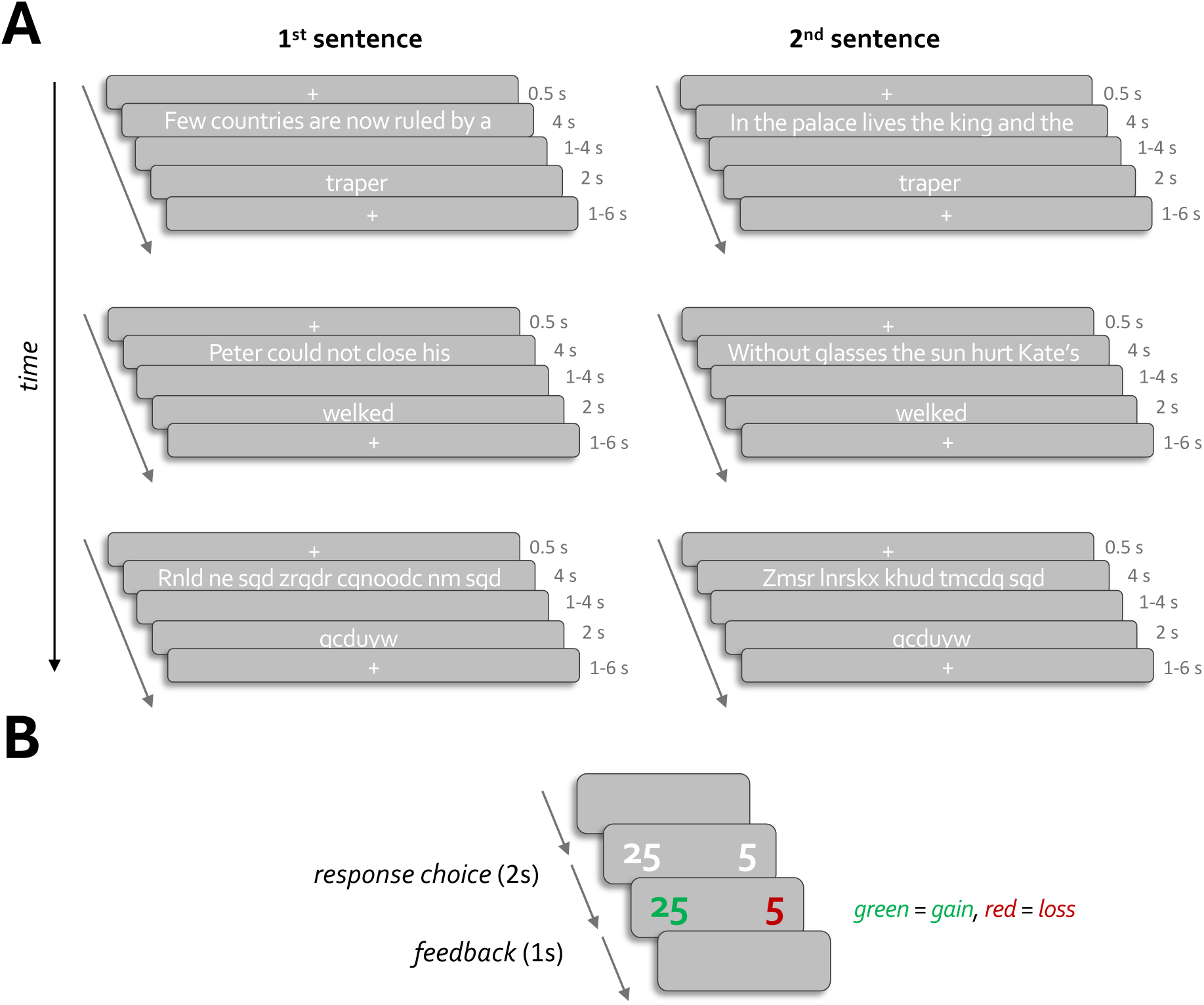
Experimental tasks: **A**) **Schematic overview of the contextual word learning task.** Participants were instructed to learn the meanings of pseudowords embedded within pairs of sentences. The first sentence provided limited contextual information, whereas the second sentence provided much more contextual information to infer the pseudoword’s meaning. Sentence pairs were presented non-consecutively. Non-readable stimuli were included to control for visual processing demands. **B**) **Schematic overview of the monetary reward task**. On each trial, participants selected one of two numbers. Following their choice, one number was highlighted in green and the other in red. A green outcome indicated a monetary gain in pence equal to the displayed value, whereas a red outcome indicated a monetary loss of that value.

The 36 pseudowords were presented across six scanning runs, with six novel pseudowords introduced per run. Following previous implementations of the paradigm (Ripollés et al., 2014, 2016, 2018), the two sentences associated with each pseudoword were presented non-consecutively. This design more closely approximates naturalistic word learning, in which encounters with unfamiliar words are distributed across time.

Within each run, stimuli were organised into three sets. Each set contained two pseudoword sentence pairs and one pair of non-readable control sentences. Participants first encountered the low-constraint sentences from each pair, followed later by the corresponding high-constraint sentences (Fig. 1A). This structure ensured that the second sentence of a pair was separated from the first by at least two intervening sentences and by no more than five intervening sentences. The order of sentences within sets, sets within runs, and runs within the experiment was randomised.

Stimuli were presented using PsychoPy (v2022.1.3). Each trial began with a fixation cross presented for 500 ms, followed by the sentence frame (e.g., “Few countries are now ruled by a”) displayed for 4 s. After a jittered interval of 1-4 s, a pseudoword completing the sentence (e.g., *traper*) was presented for 2 s. A second jittered fixation interval of 1-6 s followed before the onset of the next trial. Consequently, trial duration varied between 8.5 and 16.5 s. Participants were not required to make any motor responses during the learning phase, and no explicit feedback or external rewards were provided. Each run comprised 12 meaningful sentences and 6 non-readable control sentences and lasted approximately 4.5 min.

At the end of each scanning run, participants completed a three-alternative forced-choice (3AFC) recognition test assessing learning of the six pseudowords presented during that run. For each pseudoword, participants selected among three response options: the correct meaning, a foil corresponding to the meaning of another pseudoword encountered within the same run, and an “I don’t know” option. Approximately 30 min after the scanning session, participants completed a surprise delayed recognition test outside the scanner. The task followed the same 3AFC format as the in-scanner recognition tests and assessed memory for all 36 pseudowords. Pseudowords were presented in a randomised order. Chance performance on the recognition tests was 33%, as participants selected the correct answer from three response alternatives on each of 36 presented pseudowords.

##### 2.1.4.2 Monetary reward task design

An event-related monetary reward (“gambling”) task adapted from Ripollés et al. (2014) was used to independently localise reward-related brain regions (Fig. 1B). Participants completed a single run lasting approximately 8 min. Stimuli were presented using PsychoPy (v2022.1.3). Prior to scanning, participants completed a brief practice session and were instructed to earn as much money as possible during the task. They were informed that their winnings would be added to their voucher.

On each trial, two numbers were presented for 2 s. Participants selected one of the numbers using a button box held in their right hand (index finger = left option; middle finger = right option). Following the response, one number was highlighted in green and the other in red. If the selected number was highlighted in green, participants gained the corresponding amount (in pence); if highlighted in red, they lost that amount. Participants received a warning message if they failed to respond within the allotted time.

The task comprised 30 gain trials, 30 loss trials, 15 boost-gain trials, and 15 boost-loss trials. During boost trials, gains or losses were multiplied by a factor of five. In addition, participants completed 30 rest trials, each consisting of a 3 s fixation period.

At the end of the run, participants were shown their total earnings. Reward outcomes were predetermined such that participants could not develop a strategy to maximise earnings. Total earnings ranged from £0.50 to £5.00, although participants were not informed of this range before completing the task.

#### 2.1.5 Task procedure

Before scanning, participants received standardised verbal instructions for both the contextual word learning and monetary reward tasks and completed practice trials using stimuli not included in the main experiment. They were instructed to remain as still as possible throughout image acquisition.

During the scanning session, they first completed six runs of the contextual word learning task. Following each run, image acquisition was paused and participants completed the corresponding 3AFC recognition test. Participants then underwent structural MRI acquisition while watching a short movie. Finally, they completed one run of the monetary reward task.

Approximately 30 min after the completion of the word learning task, they completed a surprise delayed 3AFC recognition test outside the scanner. The neuropsychological assessment battery was administered either immediately before or after the scanning session.

#### 2.1.6 MRI acquisition

MRI data were acquired on a 3T Siemens Trio scanner equipped with a 32-channel head coil. Participants wore active noise-cancelling headphones (Optoacoustics OptoACTIVE II), and foam padding and inflatable cushions were used to minimise head motion.

Functional images were acquired using a multiband multi-echo T2*-weighted echo-planar imaging (EPI) sequence. Multi-echo acquisition improves signal recovery in regions susceptible to susceptibility-related signal dropout and facilitates denoising through echo-time-dependent signal decomposition (Lynch et al., 2020). Data acquired at the three echo times were combined into a single time series using the *tedana* pipeline (Kundu et al., 2017).

For the contextual word learning task, each run comprised 98-114 volumes, depending on the pseudorandomised jitter structure. Images were acquired with the following parameters: TR = 2300 ms; TE = 17.00, 42.01, and 67.02 ms; flip angle = 78°; field of view = 192 × 192 mm; multiband acceleration factor = 2; voxel size = 2.5 × 2.5 × 2.5 mm. Each volume consisted of 48 axial slices. For the monetary reward task, functional images were acquired using the same sequence parameters, yielding approximately 200 volumes per participant.

A high-resolution T_1_-weighted anatomical image was acquired using a magnetisation-prepared rapid gradient-echo (MPRAGE) sequence (1 mm isotropic voxels; TR = 2000 ms; TE = 2 ms; flip angle = 8°; field of view = 208 × 256 × 256 mm). The anatomical image was used for spatial normalisation and registration of functional data.

#### 2.1.7 fMRI pre-processing

Functional MRI data were preprocessed and analysed using the FMRI Expert Analysis Tool (FEAT, v6.0.5) within FSL (Smith et al., 2004). For each echo, nonbrain tissue was removed using the Brain Extraction Tool (BET; Smith, 2002). Following recommendations for multi-echo preprocessing (DuPre et al., 2021), volumes from the first echo were motion corrected using MCFLIRT (Jenkinson, 2002), and the resulting transformation parameters were applied to the remaining echoes using Applyxfm4D. Motion correction quality was assessed through visual inspection of each time series.

The three echoes were subsequently combined and denoised using the *tedana* pipeline (Kundu et al., 2017). Briefly, *tedana* performs optimal echo combination and multi-echo independent component analysis (ME-ICA) to identify and remove non-BOLD signal components. The preprocessing scripts are publicly available (https://osf.io/mkgcy/).

Following denoising, data were high-pass filtered (100 s cutoff) to remove low-frequency drift and spatially smoothed using a 5-mm full-width at half-maximum Gaussian kernel. Functional images were first aligned to a representative functional volume from the same session and then linearly registered to each participant’s T_1_-weighted anatomical image using FLIRT (Jenkinson, 2002; Jenkinson & Smith, 2001). Anatomical images were subsequently nonlinearly normalized to MNI-152 standard space using FNIRT (Andersson & Smith, 2008), and the resulting transformations were applied to the functional data.

#### 2.1.8 Statistical analysis

##### 2.1.8.1. Demographics and neuropsychological data

Group differences in sex distribution were assessed using a χ² test. All other demographic and neuropsychological measures were compared between groups using independent-samples *t* tests. A Bonferroni correction was applied to control the family-wise error rate across the six oral language and reading measures (0.05/6), yielding a corrected significance threshold of *p* < 0.008.

##### 2.1.8.2. fMRI: Contextual word learning

For both fMRI tasks (contextual word learning and monetary reward), event-related general linear models (GLMs) were implemented in FEAT. Hemodynamic responses were modelled using a double-gamma hemodynamic response function convolved with each event regressor.

For the contextual word learning task, trial onsets were modelled at the moment of the presentation of the pseudowords and with an amplitude of one. Readable pseudowords were classified as correct (i.e., successful learning) or incorrect (i.e., unsuccessful learning) using the test performed after each learning run. Twelve different conditions were modelled: (1) correct 1st pseudoword; (2) correct 1st sentence; (3) correct 2nd pseudoword; (4) correct 2nd sentence; (5) incorrect 1st pseudoword; (6) incorrect 1st sentence; (7) incorrect 2nd pseudoword; (8) incorrect 2nd sentence; (9) non-readable 1st nonword; (10) non-readable 1st sentence; (11) non-readable 2nd nonword; and (12) non-readable 2nd sentence.

First-level contrasts were specified for all participants using each of the 12 conditions against the baseline. These contrast images were introduced into a second-level repeated-measures analysis of variance (ANOVA) that also modelled the subject-specific constants. To confirm that the readable stimuli activated children’s reading networks, we first compared whole brain fMRI-responses for the readable sentences against the non-readable stimuli (correct 1st sentence + incorrect 1st sentence + correct 2nd sentence + incorrect 2nd sentence > non-readable 1st sentence + non-readable 2nd sentence). Our key contrast of interest was designed to delineate the difference in neural activity between the correct versus incorrect pseudowords taken at the second presentations (correct 2nd pseudoword > incorrect 2nd pseudoword). The second key contrast was the difference between the second versus first pseudoword presentations regardless of learning success (correct 2nd pseudoword + incorrect 2nd pseudoword > correct 1st pseudoword + incorrect 1st pseudoword).

We lastly performed a between-group comparison of TD and RD groups in a higher-level analysis using FMRIB’s Local Analysis of Mixed Effects (FLAME; Beckmann et al., 2003; Woolrich et al., 2004). Z (Gaussianised T/F) statistic images were thresholded using clusters determined by Z > 3.1 and a (corrected) cluster significance threshold of *p* = 0.05 (Worsley, 2001).

##### 2.1.8.3. fMRI: Monetary reward

For the monetary reward task, the GLM included four regressors corresponding to monetary gains, losses, boost gains, and boost losses. Events were modelled with a duration of 0.001 s and amplitude of one. To identify reward-responsive brain regions, the primary contrast compared rewarding versus nonrewarding outcomes (all gains > all losses), where gains included both standard and boost-gain trials and losses included both standard and boost-loss trials. The same between-group comparison of TD and RD groups described in previous section was performed for this task.

##### 2.1.8.4. fMRI: Region-of-interest analysis

To further characterise reward-related activity during word learning, we performed a region-of-interest (ROI) analysis using an independent functional localiser derived from the monetary reward task. A higher-level group analysis across all participants identified regions showing greater activation for gains than losses (all gains > all losses). Statistical maps were thresholded at Z > 3.1 with cluster-corrected *p* < 0.05. From these thresholded maps, we extracted bilateral masks corresponding to the left and right ventral striatum.

For each participant, mean parameter estimates were extracted from the left and right ventral striatal ROIs for the contextual word learning task conditions. These values were analysed in R (v4.3.2; R Core Team, 2023) using a mixed-design ANOVA with Group (TD, RD) as a between-subjects factor and Presentation Order (1st, 2nd) and Learning Accuracy (correct, incorrect) as within-subject factors. We examined the effect of all three factors as well as all interactions among them, on ventral striatal activity. Post-hoc comparisons used Tukey’s method.

We also conducted Pearson’s correlations to evaluate the relationship between children’s reading scores (as measured through TOWRE-II, see methods section 2.1.3) and ventral striatal activity extracted from the ROI during word learning. Specifically, we examined the relationship between reading scores, and successful word learning (i.e., correct 2nd pseudowords), unsuccessful word learning (i.e., incorrect 2nd pseudowords), 2nd non-readable nonwords, and lastly ventral striatal activity during monetary gains (i.e., all gains > all losses).

### 2.2. Secondary data analysis

To examine the relationship between reading ability and the subjective experience of word learning, we conducted a secondary analysis of an openly available dataset previously reported by Bains et al. (2023). The original study included 345 children and adolescents aged 11-18 years. The present analysis focused on the subset of participants aged 11-13 years (*N* = 68) to match the age range of our fMRI sample.

#### 2.2.1 Ethics

The study was approved by the Central Ethics Committee at Royal Holloway, University of London (2402-2020-11-09-20-05-UGJT014). Written parental consent and participant assent were obtained prior to participation.

#### 2.2.2. Participants

Participants were native English speakers with normal or corrected-to-normal vision. Individuals with any known neurological conditions or speech, language, or hearing disorders were excluded.

#### 2.2.3. Word learning task

Participants completed an online contextual word-learning task administered through Gorilla (www.gorilla.sc). The task consisted of 40 sentence pairs ending in novel pseudowords (Fig. 2A). In 20 congruent trials (M+), the two sentences supported a single interpretable meaning for the pseudoword. For instance, “Few countries are now ruled by a traper” followed by “In the palace lives the king and the traper” suggests that *traper* means *queen*. In the remaining 20 incongruent trials (M−), the second sentence was manipulated such that no consistent meaning could be inferred. For example, “Old people sometimes lose their jedin” implies *jedin* could mean *hair*, but the scrambled second sentence “Joan fed the baby some warm jedin” now implies that *jedin* means *milk*, making a consistent meaning impossible. Congruent and incongruent conditions were matched for novelty, task structure, and cognitive demands and were counterbalanced across stimulus lists (Mestres-Misse et al., 2007; Ripollés et al., 2014, 2016, 2018).

**Figure 2.**
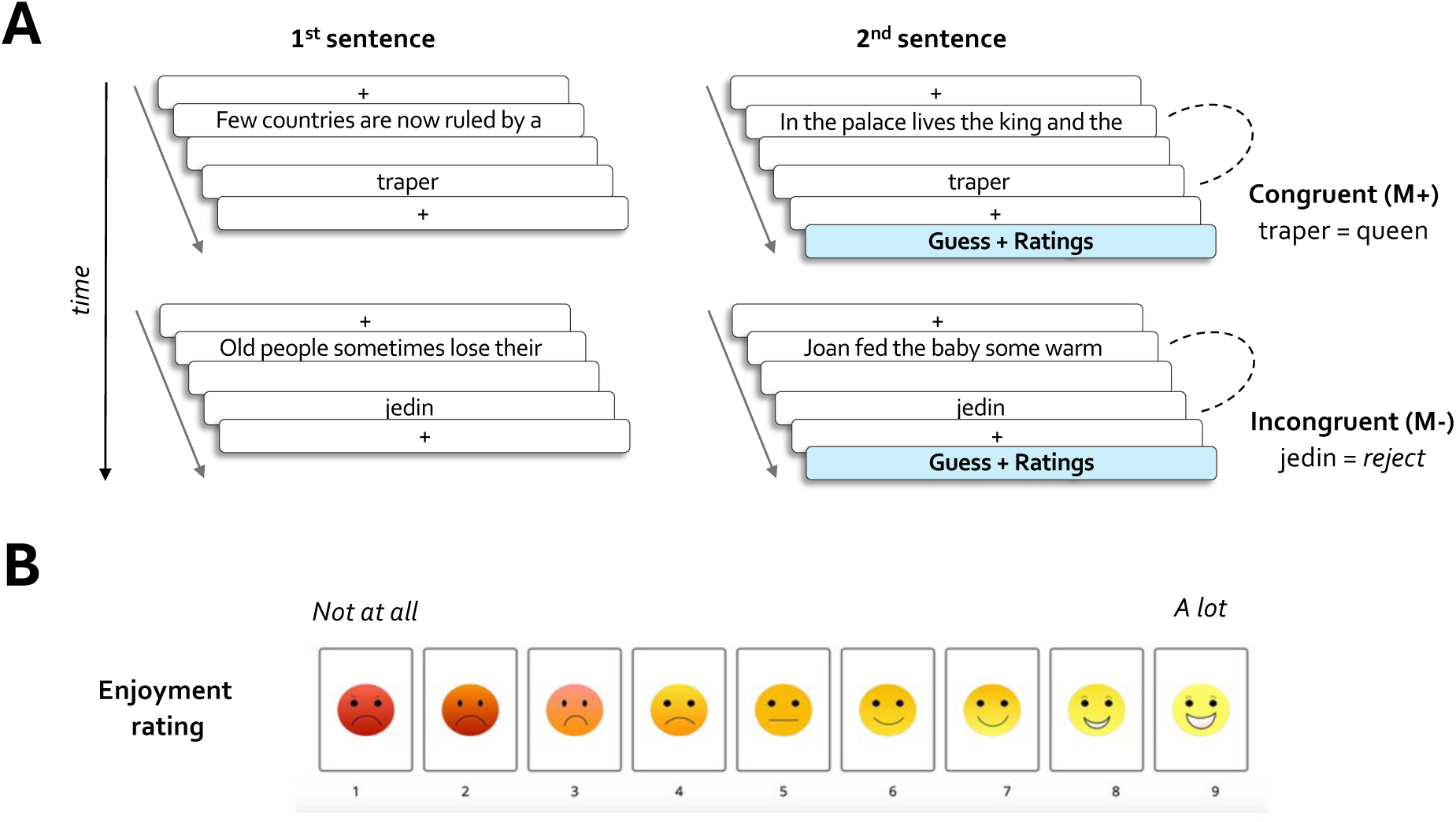
Secondary behavioural study: **A**) **Schematic overview of the contextual word learning task**. Participants encountered sentence pairs containing novel pseudowords. In the congruent condition (M+), a consistent meaning could be inferred across the two sentences. In the incongruent condition (M−), no single meaning could be derived, and participants were required to reject the pseudoword. **B**) **Enjoyment ratings collected following each word learning trial**. Participants rated how much they enjoyed learning the pseudoword using a 9-point visual scale.

After reading each sentence pair, participants either typed the inferred meaning of the pseudoword (M+ trials) or entered “reject” when they judged the pair to be incongruent (M− trials). Participants then rated how much they enjoyed learning the pseudoword using a 9-point visual scale (Fig. 2B). Participants also encountered three “catch” trials interspersed throughout the learning phase, which required choosing specific buttons to confirm attention and compliance. No feedback was provided during the task.

Participants next completed the Rapid Online Assessment of Reading Lexical Decision Task (ROAR-LDT), a lexical decision task that closely relates to reading ability (Yeatman et al., 2021). Participants were excluded if they did not complete any of the catch trials accurately, performed below 25% accuracy on congruent trials, or provided the same enjoyment rating across all trials.

### Statistical analysis

Participants were classified as stronger or poorer readers using a median split on ROAR-LDT performance. Enjoyment ratings were analysed using a 2 × 2 × 2 mixed-design ANOVA with Reading Ability (strong readers, poor readers) as a between-subjects factor and Learning Condition (congruent, incongruent) and Learning Accuracy (correct, incorrect) as within-subject factors. Post-hoc comparisons used Tukey’s method.

## 3. Results

### 3.1 Whole brain fMRI results

#### 3.1.1 Neural activity during sentence reading

We first verified that reading sentences, relative to viewing non-readable sentences, elicited fMRI activity in children in brain regions associated with reading (contrast: correct 1st sentence + incorrect 1st sentence + correct 2nd sentence + incorrect 2nd sentence > non-readable 1st sentence + non-readable 2nd sentence). As expected, we observed activity in a primarily left-lateralised network of regions in both neurotypical (TD) children and dyslexic (RD) children, which included the left inferior frontal gyrus and middle temporal gyrus, the inferior occipital cortex (including fusiform gyrus), and the lateral occipital cortex in both hemispheres (Fig. 3). Activity was also seen in the ventral striatum, insular cortex and the thalamus bilaterally. There were no significant group differences.

**Figure 3.**
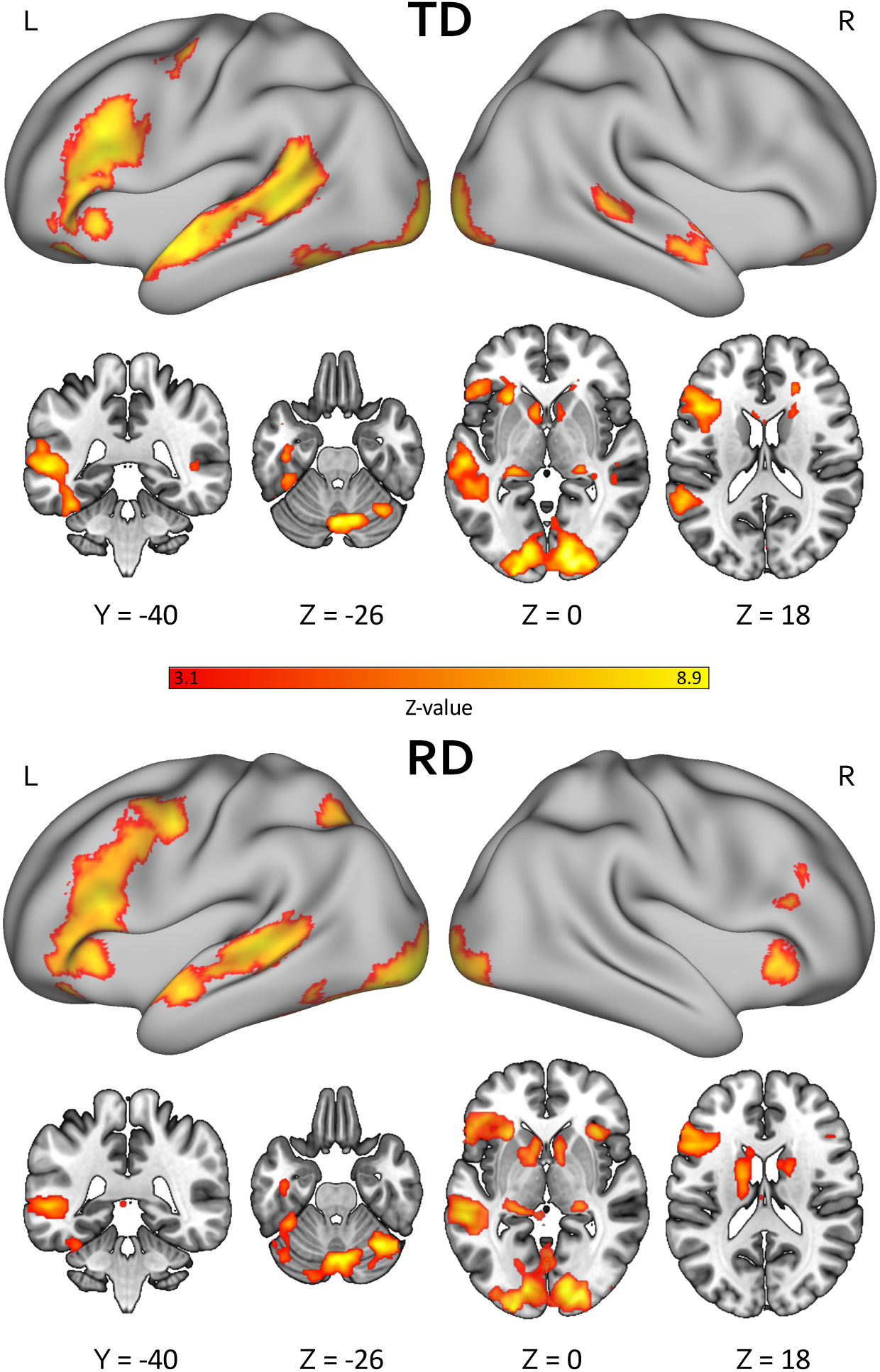
Whole-brain activation for readable relative to non-readable sentences. Statistical maps show mean fMRI activation for the contrast of readable vs. non-readable sentences (Z > 3.1, *p* < 0.05, FWE-corrected) in neurotypical (TD) and dyslexic (RD) children. Numbers below each slice indicate MNI coordinates (mm) relative to the anterior commissure. L, left; R, right.

#### 3.1.2 Increased ventral striatal activity during word learning in predictable contexts

At the whole-brain level, contrary to our predictions, we did not observe neural differences related to successful versus unsuccessful word learning (contrast: correct 2nd pseudoword > incorrect 2nd pseudoword) in either group. Given that we did not provide feedback, children may not have had the same metacognitive insight into their performance as adults (Ripollés et al., 2014). We therefore examined brain activity associated with words that appeared in the second presentation of the sentences relative to the first, regardless of learning success (i.e., correct 2nd pseudoword + incorrect 2nd pseudoword > correct 1st pseudoword + incorrect 1st pseudoword). This contrast allowed us insight into when words could be inferred. We observed increased neural activity in the ventral striatum and caudate nuclei bilaterally in the neurotypical group specifically, as well as a left-lateralised frontoparietal network, encompassing the left frontal pole, the superior frontal gyrus more medially, inferior and middle frontal gyri, left inferior parietal cortex extending into lateral occipital cortex, and the anterior insula bilaterally (Fig. 4; Table 2). We did not observe any significant differences between neurotypical and dyslexic children, though the activations were much less extensive in the dyslexic children.

**Figure 4.**
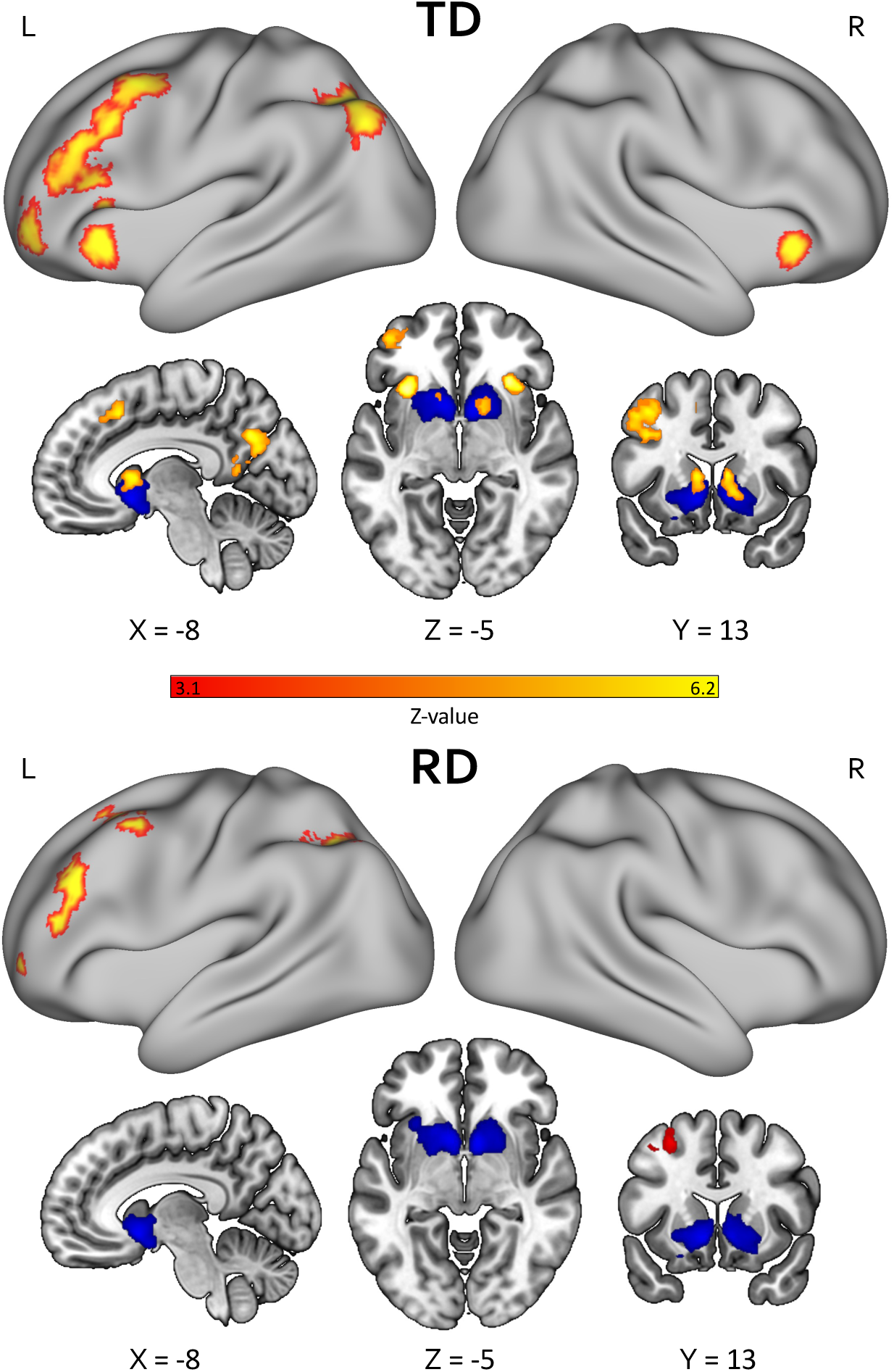
Neural responses to second vs. first pseudoword presentations, and monetary reward processing across groups. fMRI activations associated with the contrast of the second versus first pseudoword presentation, irrespective of learning success, are shown in red/yellow for neurotypical (TD) and dyslexic (RD) children. Greater activation during the second presentation was observed in ventral striatal regions and overlapped with activation elicited by monetary reward processing, defined by the contrast of all gains versus all losses in the monetary incentive task (blue masks). Statistical maps were thresholded at *Z* > 3.1, *p* < 0.05, family-wise error (FWE) corrected. No significant between-group differences were observed, although activation was less spatially extensive in the RD group. Numbers below each slice indicate MNI coordinates (mm) relative to the anterior commissure. L, left; R, right.

**Table 2.** Peak coordinates for the second versus first pseudoword presentation contrast across the whole cohort.

| Anatomical area | Coordinates |  |  | Z-statistic | Voxels | p-value |
| --- | --- | --- | --- | --- | --- | --- |
|  | X | Y | Z |  |  |  |
| R Frontal pole | 36 | 52 | 12 | 4.00 | 133 | 0.0358 |
| L Middle frontal gyrus | -42 | 8 | 50 | 5.79 | 4218 | <0.0001 |
| R Middle frontal gyrus | 32 | 10 | 44 | 4.23 | 142 | 0.0274 |
| R Insular cortex | 34 | 22 | 0 | 5.24 | 160 | 0.0163 |
| L Caudate nucleus | -8 | 12 | 6 | 4.76 | 268 | 0.0009 |
| R Caudate nucleus | 10 | 12 | 8 | 4.61 | 140 | 0.0291 |
| L Posterior middle temporal gyrus | -64 | -28 | -10 | 4.29 | 151 | 0.0211 |
| L Angular gyrus | -38 | -54 | 40 | 6.18 | 1454 | <0.0001 |
| L Precuneus cortex | -12 | -64 | 28 | 5.10 | 1133 | <0.0001 |
| R Superior lateral occipital cortex | 42 | -58 | 48 | 4.10 | 180 | 0.0092 |
| R Crus I cerebellum | 32 | -64 | -30 | 4.29 | 232 | 0.0023 |
| R Crus II cerebellum | 6 | -82 | -36 | 5.45 | 713 | <0.0001 |

*Note.* Statistical maps for the contrast of second versus first pseudoword presentations (regardless of learning success), thresholded at $Z > 3.1$ , $p < 0.05$ , family-wise error (FWE) corrected. Peak coordinates are reported for all significant clusters (see also red/yellow activations in Fig. 4). Coordinates are given in MNI space as $x$ (sagittal), $y$ (coronal), and $z$ (axial) positions in mm relative to the anterior commissure, together with peak $Z$ value and cluster extent (voxels). L, left; R, right.

#### 3.1.3 Shared ventral striatal activity across monetary reward and word learning tasks

We localised brain regions implicated in reward processing using an independent monetary reward localiser. Monetary gain (contrast: all gains > all losses) was associated with robust activation in the ventral striatum across the whole cohort, with no group differences in reward processing between TD and RD groups (Fig. S1 and blue areas in Fig. 4). Notably, both this monetary reward task and the word learning contrast, where meaning could be inferred (i.e., correct 2nd pseudoword + incorrect 2nd pseudoword > correct 1st pseudoword + incorrect 1st pseudoword) elicited activity in an overlapping region in the ventral striatum.

### 3.2 ROI analyses

#### 3.2.1 Increased ventral striatal activity during successful word learning in neurotypical but not dyslexic children

A region-of-interest (ROI) analysis was carried out to assess whether activity in the ventral striatal region sensitive to monetary reward was modulated by word learning (see methods section 2.1.8.4). A 2 × 2 × 2 mixed ANOVA ((Group: TD, RD) × (Presentation Order: 1st, 2nd) × (Learning Accuracy: correct, incorrect)) revealed a significant three-way interaction between Group, Presentation Order and Learning Accuracy [*F*(1, 43) = 4.25, *p* = 0.045, η² = 0.011; Fig. 5A]. In the successful word learning trials, the neurotypical children had significantly higher ventral striatal activity for words appearing in the second presentation of the sentences compared with the first [t(43) = -3.86, *SE* = 0.015, *p*_adj_ = 0.008]. Dyslexic children did not exhibit the same increase in activity in successful trials (*p*_adj_ = 0.910). For unsuccessful trials, this increase was not observed in either the neurotypical (*p*_adj_ = 0.957) or dyslexic children (*p*_adj_ = 0.158). There was a main effect of Presentation Order, with greater activity observed for words embedded in the 2nd sentence relative to the 1st (i.e., main effect of Presentation Order: *F*(1, 43) = 13.87, *p* < 0.001, η² = 0.049; Fig. S2). Further, the Group × Learning Accuracy interaction was significant [*F*(1, 43) = 4.39, *p* = 0.042, η² = 0.019], but post-hoc pairwise comparisons did not reveal significant differences between groups. This may have been driven by a trend for greater striatal activation during incorrect relative to correct sentences in the dyslexic children (*p*_adj_ = 0.094). No other main effect or interactions were significant.

**Figure 5.**
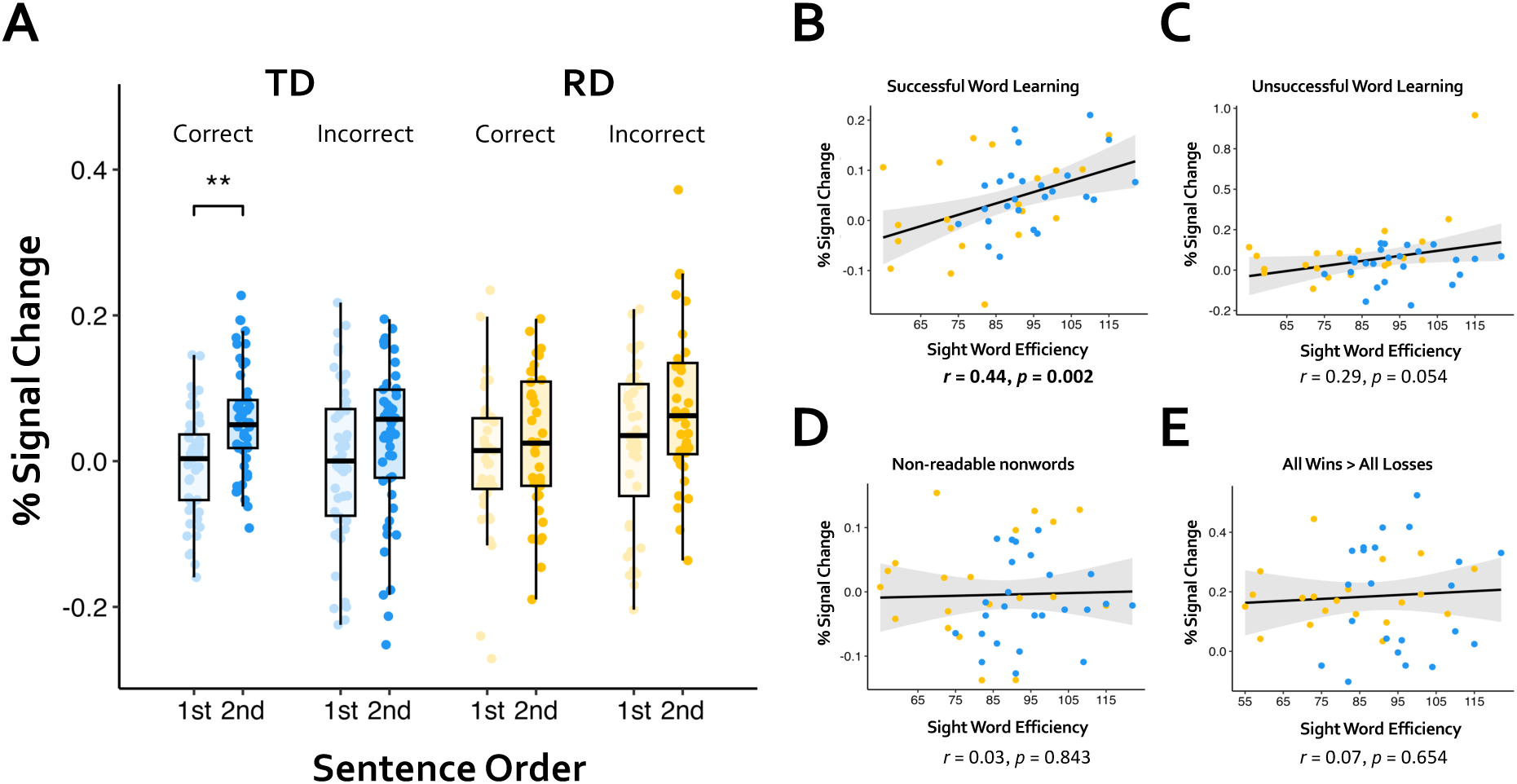
Ventral striatal activity during word learning and its association with reading ability. **(A)** Ventral striatal activity as a function of Presentation Order (2nd vs. 1st pseudoword presentation) and Learning Accuracy (correct vs. incorrect meaning inference) in neurotypical (TD, in blue) and dyslexic (RD, in yellow) children. Boxplots show the median, interquartile range (IQR), and whiskers extending to ±1.5 × IQR. Paler colours indicate first presentations, darker colors indicate second pseudoword presentations. **(B)** Association between ventral striatal activity during successful word learning (correct meaning inference) and reading ability, assessed using the Sight Word Efficiency subtest of TOWRE-2. Higher reading scores were associated with greater ventral striatal activity during successful word learning. No significant associations with reading ability were observed for **(C)** unsuccessful word learning (incorrect meaning inference), **(D)** presentations of non-readable nonwords, or **(E)** reward-related ventral striatal activity during the monetary incentive task (all gains > all losses).

#### 3.2.2 Ventral striatal activity during word learning is associated with reading ability

We also determined if activity in ventral striatum during word learning was associated with a continuous measure of reading ability, which was measured using the Sight Word Efficiency subtest from Test of Word Reading Efficiency-Second Edition (TOWRE-2; Torgesen et al., 2012). Better readers showed greater ventral striatal activity during successful word learning trials (i.e., correctly inferring the meaning of pseudowords at their second presentations: *r* = 0.44, *p* = 0.002; Fig. 5B). Importantly, this was not the case for unsuccessful word learning (i.e., incorrectly inferring the meaning of pseudowords at their second presentations: *r* = 0.29, *p* = 0.054; Fig. 5C), or non-readable nonwords (i.e., nonwords at their second presentations: *r* = 0.03, *p* = 0.843; Fig. 5D). As shown in Figure 5C, there was an outlier (one RD participant) in the correlation between unsuccessful word learning and reading ability. On removal of this outlier, this correlation was still not significant (*r* = 0.151, *p* = 0.329). To ascertain the specificity of the relationship between reading ability and reward processing, we also examined if reading ability was related to ventral striatal activity for the monetary reward task (i.e., all gains > all losses contrast). There was no association between activity in the ventral striatal ROI for monetary reward and reading ability (*r* = 0.07, *p* = 0.654; Fig. 5E).

### 3.3 Secondary data analysis

#### 3.3.1 Better readers report greater enjoyment ratings for successful word learning

To assess whether differences observed in the reward processing system were also reflected in behaviour, we conducted a secondary data analysis using an independent developmental dataset (Bains et al., 2024). (Figs. 2 and 6). From this dataset of 345 children, we elected 68 children aged 11-13 years, and divided them into strong and poor readers based on a median split of performance on an online lexical decision task reflecting reading ability (Yeatman et al., 2021). We examined behavioural ratings of enjoyment and how these varied based on learning accuracy (correct/ incorrect) in congruent (M+) and incongruent (M-) trials.

**Figure 6.**
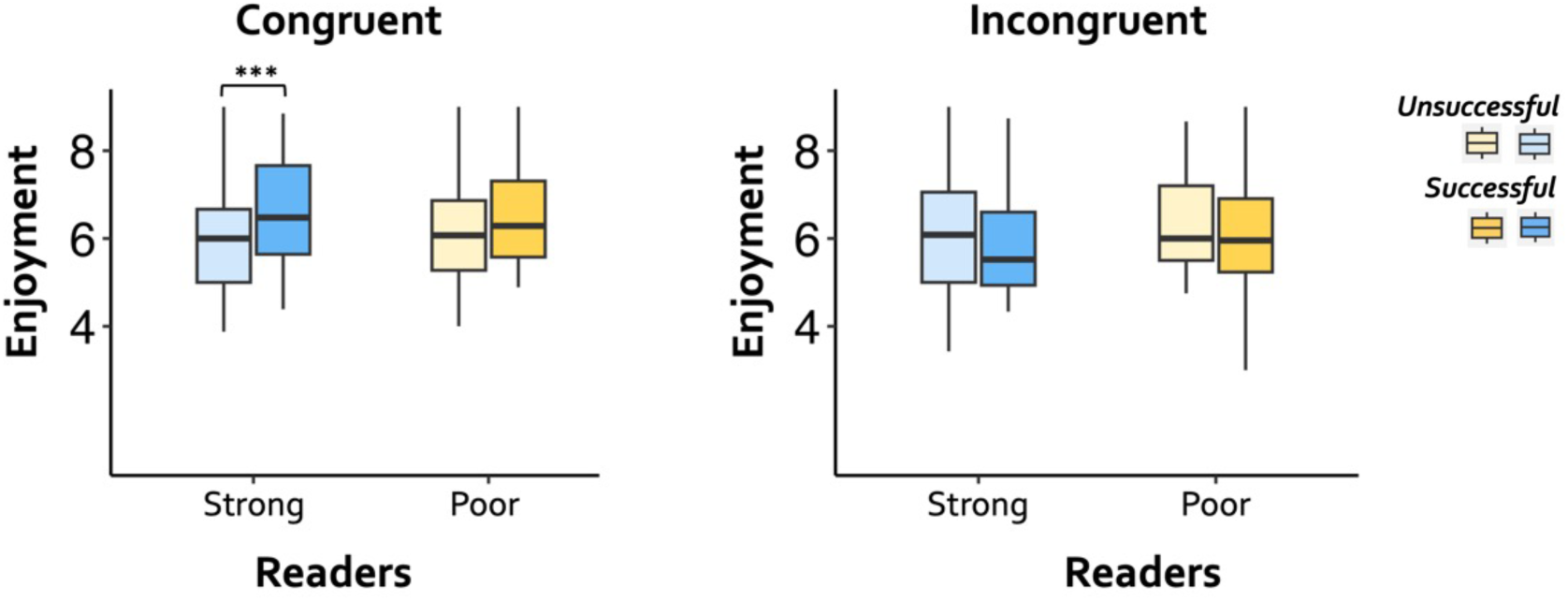
Better readers report greater enjoyment during congruent word learning. Enjoyment ratings as a function of Reading Ability (strong readers, poor readers), Learning Accuracy (correct, incorrect), and Learning Condition (congruent, incongruent). In the congruent condition, where pseudoword meaning could be inferred from context, children with higher reading ability reported greater enjoyment for successful (correct) relative to unsuccessful (incorrect) word-learning trials. This effect was not observed among poorer readers. The y-axis represents self-reported enjoyment ratings. Boxplots show the median, interquartile range (IQR), and whiskers extending to ±1.5 × IQR. Darker shades indicate successful (correct) trials, paler shades indicate unsuccessful (incorrect) trials. \*\**p* < 0.005.

The 2 × 2 × 2 mixed design ANOVA ((Reading Ability: strong readers, poor readers) × (Learning Accuracy: correct, incorrect) × (Learning Condition: congruent, incongruent)) revealed a three-way interaction between these factors [F(1, 62) = 7.66, *p* = 0.007, η² = 0.004]. Post hoc testing revealed that strong readers showed greater enjoyment for successful word learning, and this effect was modulated by congruency. In other words, strong readers showed greater enjoyment only in congruent trials and when they were successful [t(62) = -5.87, *SE* = 0.114, *p*_adj_ = 0.001]. Strong readers did not exhibit enhanced enjoyment during incongruent word learning trials regardless of success (*p*_adj_ = 0.091). In contrast, poor readers did not report significantly greater enjoyment when successfully learning words in the congruent condition (*p*_adj_ = 0.785), or during incongruent trials (*p*_adj_ = 0.995), further reinforcing the idea that children who are better at monitoring and evaluating success experience greater reward during word learning.

## 4. Discussion

We predicted that children would generate intrinsic reward activity when successfully learning new words. Consistent with adult findings, we observed neural activity in reward processing regions when children made predictions about the meaning of novel words (Ripollés et al., 2014, 2016). We also explored how reading proficiency might modulate this intrinsic response. We found reading ability exerted a strong influence on the degree of intrinsic reward experienced by children during word learning, with children with dyslexia showing reduced activity in the ventral striatum, even when they successfully learned words. This was mirrored in our analyses of behavioural data, with reading ability associated with lower enjoyment of word learning, even when learning was successful. These findings provide a better mechanistic understanding for why reading may be less enjoyable in dyslexic learners.

Our findings (particularly in neurotypical children) clearly indicate that the generation and subsequent confirmation of linguistic predictions about word meaning is associated with increased activity in reward-processing regions. This is the first demonstration of this effect in children at the neural level, converging with behavioural effects in this domain (Bains et al., 2024). One reason we may have been uniquely sensitive to striatal responses is our use of a multi-echo sequence that minimised signal drop-out in the striatum. Our findings align with adult studies showing that predictive language learning is associated with activity in the ventral striatum (Orpella et al., 2021; Ripollés et al., 2014), behavioural ratings of pleasure (Zaka et al., 2026), and galvanic skin responses (Ripollés et al., 2016). Importantly, our results indicate that these intrinsic responses characterise learning across the lifespan, echoing their putative evolutionary importance (Syal & Finlay, 2011). Previous work has shown that such affective responses can drive later learning and engagement, with reward linked to improved memory during word learning (Garvin & Krishnan, 2022; Zaka et al., 2026) as well as in other domains such as semantic learning (Gruber et al., 2014). There is wide variability in leisure reading, with readers in the 90% percentile encountering ∼1.8 million words per year, and readers in the 10% percentile encountering about 8000 (Anderson et al., 1988). Particularly given recent declines in reading for pleasure (Clark et al., 2023), these affective responses are critical to understand.

Interestingly, individual differences in reading ability were a crucial factor influencing reward processing during word learning. Only strong readers exhibited greater ventral striatal activity during successful word learning. This association was specific to the successful word learning trials, as this uptick in striatal activity was not observed in unsuccessful word learning trials or in non-readable trials. Critically, this association could not be attributed to a general alteration of reward processing, as there were no significant group differences in ventral striatal activity for monetary reward. To be confident that our interpretation of the fMRI data was not reverse inference, we supplemented the imaging results with a secondary data analysis of behavioural enjoyment ratings in word learning. Our behavioural results mirrored the findings observed at the brain level, showing that stronger readers rated successful word learning in trials, wherein correct meaning acquisition was possible (congruent condition), as more enjoyable. Further, they did not show increased enjoyment for incongruent sentences, where it is possible to be accurate in one’s performance in the task without acquiring meaning. While population-based studies have suggested that reading ability might be a driver of enjoyment, our study is the first to show the neural mechanisms by which this drive might occur (Bridges, 2014; Froiland & Oros, 2014; Van Bergen et al., 2018)

We speculate that metacognitive insight into performance might drive the generation of intrinsic reward. Children did not receive explicit feedback on the accuracy of their performance in the word learning paradigm. Intrinsic reward, especially during contextual word learning, is highly related to performance monitoring or confidence in one’s ability (Ripollés et al., 2016; Vavra et al., 2023), which in this context translates to the ability to track one’s learning of the linguistic material. These previous results in adults suggest that better readers can draw on their knowledge and experience to benefit from the confirmation of a prediction about meaning; that is, from successful word learning. In contrast, our dyslexic participants might have simply been less confident about their predictions, thereby experiencing diminished intrinsic reward even when they were successful in learning the novel words.

These results have clear practical implications. Our findings suggest that even when poor readers succeed in a task, they might not experience the same level of reward as their peers. This diminished sense of reward may contribute to their reluctance to engage in reading. Therefore, simply increasing exposure to reading is unlikely to change their lowered intrinsic motivation. Perhaps a more effective strategy would be to create a “shift” in the reader’s motivational state by boosting self-efficacy. For instance, explicitly asking the child to generate predictions and providing feedback to feel more confident about their predictions might make the interactions more rewarding. This approach aligns well with research demonstrating that making interventions more motivational lead to better outcomes (McBreen & Savage, 2022). It is also consistent with previous work on motivation and curiosity showing that learning is enhanced when learners are motivated to resolve gaps in their knowledge and when outcomes exceed expectations (Garvin & Krishnan, 2022). From this perspective, interventions that increase curiosity, prediction generation, and the rewarding experience of successful comprehension may not only improve engagement with reading but also strengthen learning and memory.

A limitation of our work is that our conclusions are based on a relatively small sample of children. However, while replication is critical, it is important to note this was a focused exploration of striatal signals in word learning using a paradigm that has been well studied in adults, and more recently, behaviourally in children. We were able to collect 36 minutes of task imaging data from a well-characterised cohort. Critically, we scanned children with a range of reading abilities, allowing us to study a broader distribution.

In conclusion, this study points to an important role for reward systems in word learning during development. The ventral striatum, a core region associated with reward, is activated when children generate and confirm predictions about word meanings. The generation of this signal is modulated by reading skills, with better readers showing heightened enjoyment and increased brain activity in reward-related regions during successful word learning. This provides a mechanistic explanation for how reading ability may drive a virtuous cycle of learning in children and adolescents and provides clues into targets for intervention.

## 5. Data and code availability

All statistical analyses were conducted using the R programming language (R Core Team, 2022). The anonymised neuropsychological scores and codes supporting the findings of this study are openly available on the Open Science Framework (https://osf.io/d8ra3).

## 7. Acknowledgements

We thank all the children and families who participated in this study; this work would not have been possible without their time and commitment. We gratefully acknowledge financial support from the Experimental Psychology Society, the Academy of Medical Sciences (SBF006/1031) and the UKRI Medical Research Council (MR/X003647/2).

## References

1. Anderson, R. C., Wilson, P. T., & Fielding, L. G. (1988). Growth in Reading and How Children Spend Their Time outside of School. Reading Research Quarterly. http://eric.ed.gov/?id=EJ373263

2. Andersson, J., & Smith, S. (2008). FNIRT-FMRIB’s non-linear image registration tool. Human Brain Mapping.

3. Angwin, A. J., Wilson, W. J., Ripollés, P., Rodriguez-Fornells, A., Arnott, W. L., Barry, R. J., Cheng, B. B. Y., Garden, K., & Copland, D. A. (2019). White noise facilitates new-word learning from context. Brain and Language, 199, 104699. 10.1016/j.bandl.2019.104699

4. Bains, A., Barber, A., Nell, T., Ripollés, P., & Krishnan, S. (2024). The role of intrinsic reward in adolescent word learning. *Developmental Science*, e13513. 10.1111/desc.13513

5. Beckmann, C. F., Jenkinson, M., & Smith, S. M. (2003). General multilevel linear modeling for group analysis in FMRI. NeuroImage, 20(2), 1052–1063. 10.1016/S1053-8119(03)00435-X

6. Bridges, L. (2014). The Joy and Power of Reading. Scholastic Inc.

7. Bromberg-Martin, E. S., Matsumoto, M., & Hikosaka, O. (2010). Dopamine in Motivational Control: Rewarding, Aversive, and Alerting. Neuron, 68(5), 815–834. 10.1016/j.neuron.2010.11.022

8. Caras, M. L., Happel, M. F. K., Chandrasekaran, B., Ripollés, P., Keesom, S. M., Hurley, L. M., Remage-Healey, L., Holt, L. L., & Wright, B. A. (2022). Non-sensory Influences on Auditory Learning and Plasticity. Journal of the Association for Research in Otolaryngology, 23(2), 151–166. 10.1007/s10162-022-00837-3

9. Clark, C., Bonafede, F., Picton, I., & Cole, A. (2023). Children and young people’s writing in 2023. London: National Literacy Trust.

10. Decker, A. L., Meisler, S. L., Hubbard, N. A., Bauer, C. C. C., Leonard, J., Grotzinger, H., Giebler, M. A., Torres, Y. C., Imhof, A., Romeo, R., & Gabrieli, J. D. E. (2024). Striatal and Behavioral Responses to Reward Vary by Socioeconomic Status in Adolescents. The Journal of Neuroscience, 44(11), e1633232023. 10.1523/JNEUROSCI.1633-23.2023

11. DuPre, E., Salo, T., Ahmed, Z., Bandettini, P., Bottenhorn, K., Caballero-Gaudes, C., Dowdle, L., Gonzalez-Castillo, J., Heunis, S., Kundu, P., Laird, A., Markello, R., Markiewicz, C., Moia, S., Staden, I., Teves, J., Uruñuela, E., Vaziri-Pashkam, M., Whitaker, K., & Handwerker, D. (2021). TE-dependent analysis of multi-echo fMRI with tedana. Journal of Open Source Software, 6(66), 3669. 10.21105/joss.03669

12. Froiland, J. M., & Oros, E. (2014). Intrinsic motivation, perceived competence and classroom engagement as longitudinal predictors of adolescent reading achievement. Educational Psychology, 34(2), 119–132. 10.1080/01443410.2013.822964

13. Gambrell, L. B., Palmer, B. M., Codling, R. M., & Mazzoni, S. A. (1996). *Motivation to Read Profile* [Dataset]. American Psychological Association (APA). 10.1037/t43709-000

14. Garvin, B., & Krishnan, S. (2022). Curiosity-driven learning in adults with and without dyslexia. Quarterly Journal of Experimental Psychology, 75(1), 156–168. 10.1177/17470218211037474

15. Goodman, R. (1997). Strengths and Difficulties Questionnaire [Dataset]. 10.1037/t00540-000

16. Gruber, M. J., & Ranganath, C. (2019). How Curiosity Enhances Hippocampus-Dependent Memory: The Prediction, Appraisal, Curiosity, and Exploration (PACE) Framework. Trends in Cognitive Sciences, 23(12), 1014–1025. 10.1016/j.tics.2019.10.003

17. Hewitt, S. R. C., Habicht, J., Bowler, A., Lockwood, P. L., & Hauser, T. U. (2023). Probing apathy in children and adolescents with the Apathy Motivation Index–Child version. Behavior Research Methods, 56(4), 3982–3994. 10.3758/s13428-023-02184-4

18. Jenkinson, M. (2002). Improved Optimization for the Robust and Accurate Linear Registration and Motion Correction of Brain Images. NeuroImage, 17(2), 825–841. 10.1016/s1053-8119(02)91132-8

19. Jenkinson, M., & Smith, S. (2001). A global optimisation method for robust affine registration of brain images. Medical Image Analysis, 5(2), 143–156. 10.1016/s1361-8415(01)00036-6

20. Jones, H., Bains, A., Randall, L., Spaulding, C., Ricketts, J., & Krishnan, S. (2025). Investigating Reading Enjoyment in Adults With Dyslexia. Dyslexia, 31(1), e1803. 10.1002/dys.1803

21. Kuhn, M. R., & Stahl, S. A. (1998). Teaching Children to Learn Word Meanings from Context: A Synthesis and Some Questions. Journal of Literacy Research, 30(1), 119–138. 10.1080/10862969809547983

22. Kundu, P., Voon, V., Balchandani, P., Lombardo, M. V., Poser, B. A., & Bandettini, P. A. (2017). Multi-echo fMRI: A review of applications in fMRI denoising and analysis of BOLD signals. NeuroImage, 154, 59–80. 10.1016/j.neuroimage.2017.03.033

23. Lisman, J. E., & Grace, A. A. (2005). The Hippocampal-VTA Loop: Controlling the Entry of Information into Long-Term Memory. Neuron, 46(5), 703–713. 10.1016/j.neuron.2005.05.002

24. Lloyd, A., McKay, R. T., & Furl, N. (2022). Individuals with adverse childhood experiences explore less and underweight reward feedback. Proceedings of the National Academy of Sciences, 119(4), e2109373119. 10.1073/pnas.2109373119

25. Lynch, C. J., Power, J. D., Scult, M. A., Dubin, M., Gunning, F. M., & Liston, C. (2020). Rapid Precision Functional Mapping of Individuals Using Multi-Echo fMRI. Cell Reports, 33(12), 108540. 10.1016/j.celrep.2020.108540

26. Martin, N. A., & Brownell, M. A. (2011b). ROWPVT-4 Kit Receptive One-Word Picture Vocabulary Test -4. Academic Therapy Publications.

27. McBreen, M., & Savage, R. (2022). The Impact of a Cognitive and Motivational Reading Intervention on the Reading Achievement and Motivation of Students At-Risk for Reading Difficulties. Learning Disability Quarterly, 45(3), 199–211. 10.1177/0731948720958128

28. Mestres-Misse, A., Rodriguez-Fornells, A., & Munte, T. F. (2007). Watching the Brain during Meaning Acquisition. Cerebral Cortex, 17(8), 1858–1866. 10.1093/cercor/bhl094

29. Nagy, W. E., Herman, P. A., & Anderson, R. C. (1985). Learning Words from Context. Reading Research Quarterly, 20(2), 233. 10.2307/747758

30. Nation, K. (2017). Nurturing a lexical legacy: Reading experience is critical for the development of word reading skill. Npj Science of Learning, 2(1), 3. 10.1038/s41539-017-0004-7

31. Orpella, J., Mas-Herrero, E., Ripollés, P., Marco-Pallarés, J., & De Diego-Balaguer, R. (2021). Language statistical learning responds to reinforcement learning principles rooted in the striatum. PLOS Biology, 19(9), e3001119. 10.1371/journal.pbio.3001119

32. R Core Team. (2023). R: A Language and Environment for Statistical Computing. R Foundation for Statistical Computing.

33. Ripollés, P., Ferreri, L., Mas-Herrero, E., Alicart, H., Gómez-Andrés, A., Marco-Pallares, J., Antonijoan, R. M., Noesselt, T., Valle, M., Riba, J., & Rodriguez-Fornells, A. (2018). Intrinsically regulated learning is modulated by synaptic dopamine signaling. eLife, 7, e38113. 10.7554/eLife.38113

34. Ripollés, P., Marco-Pallarés, J., Alicart, H., Tempelmann, C., Rodríguez-Fornells, A., & Noesselt, T. (2016). Intrinsic monitoring of learning success facilitates memory encoding via the activation of the SN/VTA-Hippocampal loop. eLife, 5, e17441. 10.7554/eLife.17441

35. Ripollés, P., Marco-Pallarés, J., Hielscher, U., Mestres-Missé, A., Tempelmann, C., Heinze, H.-J., Rodríguez-Fornells, A., & Noesselt, T. (2014). The role of reward in word learning and its implications for language acquisition. Current Biology: CB, 24(21), 2606–2611. 10.1016/j.cub.2014.09.044

36. Rutter, Bailey, & Lord,. (2003). The Social Communication Questionnaire: Manual. Western Psychological Services, Los Angeles.

37. Siegel, L. S. (2006). Perspectives on dyslexia. Paediatrics & Child Health, 11(9), 581–587. 10.1093/pch/11.9.581

38. Smith, S. M. (2002). Fast robust automated brain extraction. Human Brain Mapping, 17(3), 143–155. 10.1002/hbm.10062

39. Smith, S. M., Jenkinson, M., Woolrich, M. W., Beckmann, C. F., Behrens, T. E. J., Johansen-Berg, H., Bannister, P. R., De Luca, M., Drobnjak, I., Flitney, D. E., Niazy, R. K., Saunders, J., Vickers, J., Zhang, Y., De Stefano, N., Brady, J. M., & Matthews, P. M. (2004). Advances in functional and structural MR image analysis and implementation as FSL. NeuroImage, 23, S208–S219. 10.1016/j.neuroimage.2004.07.051

40. Snowling, M. J., Stothard, S. E., Clarke, P., Bowyer-Crane, C., Harrington, A., Truelove, E., & Hulme, C. (2009). York Assessment of Reading for Comprehension. GL Assessment.

41. Stanovich, K. E. (1986). Matthew Effects in Reading: Some Consequences of Individual Differences in the Acquisition of Literacy. Reading Research Quarterly, 21(4), 360– 407. 10.1598/RRQ.21.4.1

42. Swanborn, M. S. L., & De Glopper, K. (1999). Incidental Word Learning While Reading: A Meta-Analysis. Review of Educational Research, 69(3), 261–285. 10.3102/00346543069003261

43. Syal, S., & Finlay, B. L. (2011). Thinking outside the cortex: Social motivation in the evolution and development of language. Developmental Science, 14(2), 417–430. 10.1111/j.1467-7687.2010.00997.x

44. Torgesen, J. K., Wagner, R. K., & Rashotte, C. A. (2012). TOWRE-2 Test of Word Reading Efficiency. Pearson.

45. Van Bergen, E., Snowling, M. J., De Zeeuw, E. L., Van Beijsterveldt, C. E. M., Dolan, C. V., & Boomsma, D. I. (2018). Why do children read more? The influence of reading ability on voluntary reading practices. Journal of Child Psychology and Psychiatry, 59(11), 1205–1214. 10.1111/jcpp.12910

46. Vavra, P., Sokolovič, L., Porcu, E., Ripollés, P., Rodriguez-Fornells, A., & Noesselt, T. (2023). Entering into a self-regulated learning mode prevents detrimental effects of feedback removal on memory. Npj Science of Learning, 8(1). 10.1038/s41539-022-00150-x

47. Wechsler, D. (2004). Wechsler Intelligence Scale for Children – Fourth UK Edition (WISC-IV UK) (L. A. Marshall & I. Coyne, Eds.; p. bpstest.2004.wisc4). British Psychological Society. 10.53841/bpstest.2004.wisc4

48. Wiig, E. H., Semel, E., & Secord, W. A. (2013). Clinical Evaluation of Language Fundamentals—Fifth Edition. Pearson Psychcorp.

49. Wittmann, M. K., Kolling, N., Akaishi, R., Chau, B. K. H., Brown, J. W., Nelissen, N., & Rushworth, M. F. S. (2016). Predictive decision making driven by multiple time-linked reward representations in the anterior cingulate cortex. Nature Communications, 7(1), 12327. 10.1038/ncomms12327

50. Woolrich, M., Behrens, T. E. J., Beckmann, C. F., Jenkinson, M., & Smith, S. M. (2004). Multi-level linear modelling for FMRI group analysis using Bayesian inference. NeuroImage, 21(4), 1732–1747.

51. Worsley, K. J. (2001). Statistical analysis of activation images. In P. Jezzard, P. M. Matthews, & S. M. Smith (Eds.), Functional Magnetic Resonance Imaging (1st ed., pp. 251–270). Oxford University PressOxford. 10.1093/acprof:oso/9780192630711.003.0014

52. Yeatman, J. D., Tang, K. A., Donnelly, P. M., Yablonski, M., Ramamurthy, M., Karipidis, I. I., Caffarra, S., Takada, M. E., Kanopka, K., Ben-Shachar, M., & Domingue, B. W. (2021). Rapid online assessment of reading ability. Scientific Reports, 11(1), 6396. 10.1038/s41598-021-85907-x

53. Zaka, H., Evans, S., Ripollés, P., & Krishnan, S. (2026). Intrinsic Reward Modulates Word Learning in Both Oral and Written Contexts. Journal of Cognition, 9(1), 28. 10.5334/joc.499

