## Supplemental Material for "Striatal activity during contextual word learning is influenced by children’s reading ability"

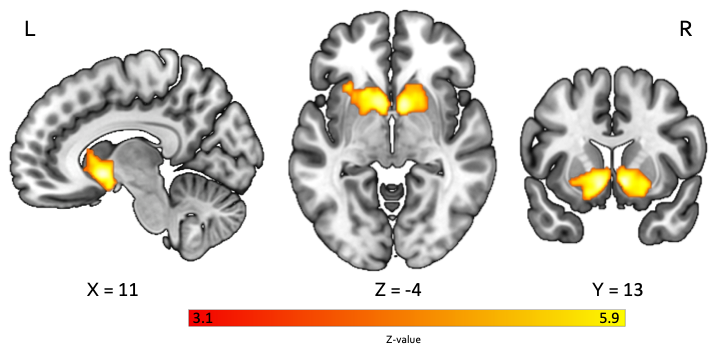

**Figure S1 (related to Fig. 4).** Brain activity associated with monetary reward across all children (n = 45). The contrast of all wins > all losses revealed significantly greater activity in the ventral striatum bilaterally (Z > 3.1, p < .05, family-wise error [FWE] corrected). Numbers beneath each slice indicate MNI coordinates (mm) relative to the horizontal and vertical planes passing through the anterior commissure. No significant differences were observed between the TD and RD groups. TD = typically developing; RD = reading difficulty; L = left; R = right.

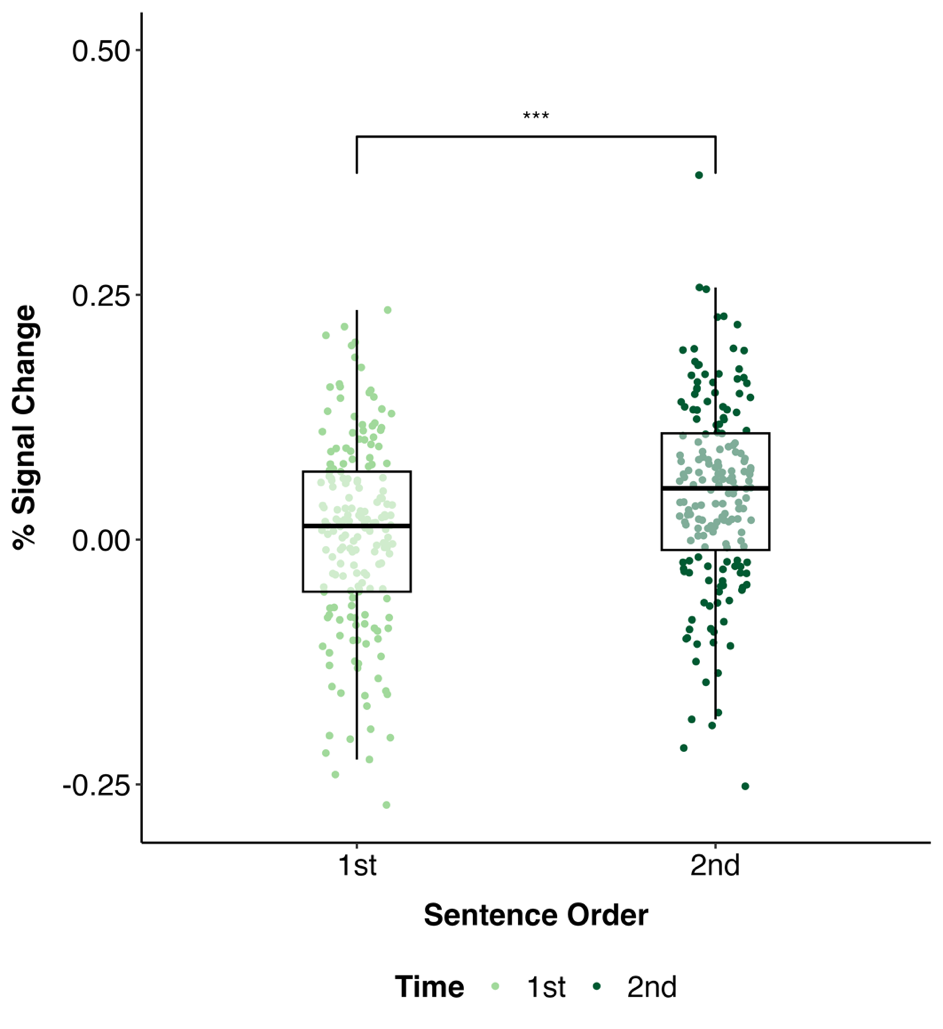

**Figure S2 (related to Fig. 5).** Enhanced ventral striatal activity during the second relative to the first presentation of words across all children (n = 45). Shaded boxes indicate the interquartile range (25th–75th percentiles), with the horizontal line denoting the median. Ventral striatal activity was significantly modulated by presentation order, with greater activity elicited by words embedded in the second sentence compared with the first (main effect of Presentation Order: F(1, 43) = 13.87, p < .001, η² = .049).
